# ActDES – a Curated Actinobacterial Database for Evolutionary Studies

**DOI:** 10.1101/2020.05.20.105536

**Authors:** Jana K. Schniete, Nelly Selem-Mojica, Anna S. Birke, Pablo Cruz-Morales, Iain S. Hunter, Francisco Barona-Gómez, Paul A. Hoskisson

## Abstract

*Actinobacteria* are a large and diverse phylum of bacteria that contains medically and ecologically relevant organisms. Many members are valuable sources of bioactive natural products and chemical precursors that are exploited in the clinic. These are made using the enzyme pathways encoded in their complex genomes. Whilst the number of sequenced genomes has increased rapidly in the last twenty years, the large size and complexity of many Actinobacterial genomes means that the sequences remain incomplete and consist of large numbers of contigs with poor annotation, which hinders large scale comparative genomics and evolutionary studies. To enable greater understanding and exploitation of Actinobacterial genomes, specialist genomic databases must be linked to high-quality genome sequences. Here we provide a curated database of 612 high-quality actinobacterial genomes from 80 genera, chosen to represent a broad phylogenetic group with equivalent genome reannotation. Utilising this database will provide researchers with a framework for evolutionary and metabolic studies, to enable a foundation for genome and metabolic engineering, to facilitate discovery of novel bioactive therapeutics and studies on gene family evolution.

**Significance as a bioresource to the community:** The *Actinobacteria* are a large diverse phylum of bacteria, often with large, complex genomes with a high G+C content. Sequence databases have great variation in the quality of sequences, equivalence of annotation and phylogenetic representation, which makes it challenging to undertake evolutionary and phylogenetic studies. To address this, we have assembled a curated, taxa-specific, non-redundant database to aid detailed comparative analysis of Actinobacteria. ActDES constitutes a novel resource for the community of Actinobacterial researchers that will be useful primarily for two types of analyses: (i) comparative genomic studies – facilitated by reliable identification of orthologs across a set of defined, phylogenetically-representative genomes, and (ii) phylogenomic studies which will be improved by identification of gene subsets at specified taxonomic level. These analyses can then act as a springboard for the studies of the evolution of virulence genes, the evolution of metabolism and identification of targets for metabolic engineering.

**Data summary:** All genome sequences used in this study can be found in the NCBI taxonomy browser https://www.ncbi.nlm.nih.gov/Taxonomy/Browser/www.tax.cgi and are summarised along with Accession numbers in Table S1

All other data are available on Figshare https://doi.org/10.6084/m9.figshare.12167529 and https://doi.org/10.5281/zenodo.3830391

a. Perl script files available on GitHub https://github.com/nselem/ActDES including details of how to batch annotate genomes in RAST from the terminal https://github.com/nselem/myrast
b. **Supp. Table S1** List of genomes from NCBI (Actinobacteria database.xlsx) https://doi.org/10.6084/m9.figshare.12167529
c. CVS genome annotation files including the FASTA files of nucleotide and amino acids sequences (individual .cvs files) https://doi.org/10.6084/m9.figshare.12167880
d. BLAST nucleotide database (.fasta file) https://doi.org/10.6084/m9.figshare.12167724
e. BLAST protein database (.fasta file) https://doi.org/10.6084/m9.figshare.12167724
f. Supp. Table S2 Expansion table genus level (Expansion table.xlsx Tab Genus level) https://doi.org/10.6084/m9.figshare.12167529
g. Supp. Table S2 Expansion table species level (Expansion table.xlsx Tab species level) https://doi.org/10.6084/m9.figshare.12167529
h. All GlcP and Glk data – blast hits from ActDES database, MUSCLE Alignment files and .nwk tree files can be found at https://doi.org/10.6084/m9.figshare.12167529
i. Interactive trees in Microreact for Glk tree https://microreact.org/project/w_KDfn1xA/90e6759e and associated files can be found at https://doi.org/10.6084/m9.figshare.12326441.v1
j. Interactive trees in Microreact for GlcP tree https://microreact.org/project/VBUdiQ5_k/0fc4622b and associated files can be found at https://doi.org/10.6084/m9.figshare.12326441.v1

## Introduction

The increase in availability of bacterial whole genome sequencing (WGS) provides large amounts of data for evolutionary and phylogenetic analysis. However, there is great variation in the quality, annotation and phylogenetic skew of the data available in large universal databases meaning that evolutionary and phylogenetic studies can be challenging. To address this, curated, high-level, taxa-specific, non-redundant sub-databases need to be assembled to aid detailed analysis. Given that there is a direct correlation between phylogenetic distance and the discovery of novel function [1–3], it is imperative that any derived databases must be phylogenetically representative and non-redundant to enable insight into the evolution of genes, proteins and pathways within a given group of taxa [1] The phylum *Actinobacteria* is a major taxon amongst the *Bacteria*, which includes phenotypically and morphologically diverse organisms found on every continent and in virtually every ecological niche [4]. They are particularly common in soils, yet within their ranks are human and animal pathogens such as *Corynebacterium, Mycobacterium, Nocardia* and *Tropheryma*, inhabitants of the gastrointestinal tract (*Bifidobacterium* and *Scardovia*) as well as plant commensals and pathogens such as *Frankia, Leifsonia* and *Clavibacter* [4, 5]. Perhaps the most notable trait of the phylum is the renowned ability to produce bioactive natural products such as antibiotics, anti-cancer agents and immuno-suppressive agents, with genera such as *Amycolatopsis, Micromonospora* and *Streptomyces* being particularly prominent [6]. As a result, computational ‘mining’ of Actinobacterial genomes has become an important part of the drug discovery pipeline, with increasing numbers of online resources and software devoted to identification of natural product biosynthetic gene clusters (BGCs)[7–9]. It is important to move beyond approaches that rely on similarity searches of known BGCs and to expand searches to identify hidden chemical diversity within the genomes [6, 7, 10–13].

A recent study of 830 Actinobacterial genomes found >11,000 BGCs comprising 4,122 chemical families, indicating that there is a vast diversity of strains and chemistry to exploit [14], yet within each of these strains there will be hidden diversity in the form of cryptic BGCs. To exploit this undiscovered diversity as the technology develops and databases expand, new biosynthetic logic will emerge, yet we know little of how natural selection shapes the evolution of BGCs and how biosynthetic precursors are supplied to them from primary metabolism and to identify targets for metabolic engineering of industrially relevant strains. These approaches will expedite industrial strain improvement processes, enabling titre increases and development of novel molecules, as well as the engineering of strains to use more sustainable feedstocks.

To aid this process we have created an Actinobacterial metabolism database including functional annotations for enzymes from 612 species to enable phylum-wide interrogation of gene expansion events that may indicate adaptive evolution, help shape metabolic robustness for antibiotic production [15] or enable the identification of targets for metabolic engineering. To demonstrate the utility of ActDES, we have detailed its construction and used it to investigate the glucose permease/glucokinase system phylogeny across the Actinobacteria.

## Methods

We generated ActDES, a database for evolutionary analysis of Actinobacterial genomes, in two ways – a database for interrogation by BLASTn or BLASTp for phylogenetic analysis and a primary metabolic gene expansion table, which can be mined at different taxonomic levels (**Supp. Table S1 and S2**) for specific metabolic functions from primary metabolism. A schematic overview of generation of the dataset is shown in **Fig.1**.

**Fig. 1.**
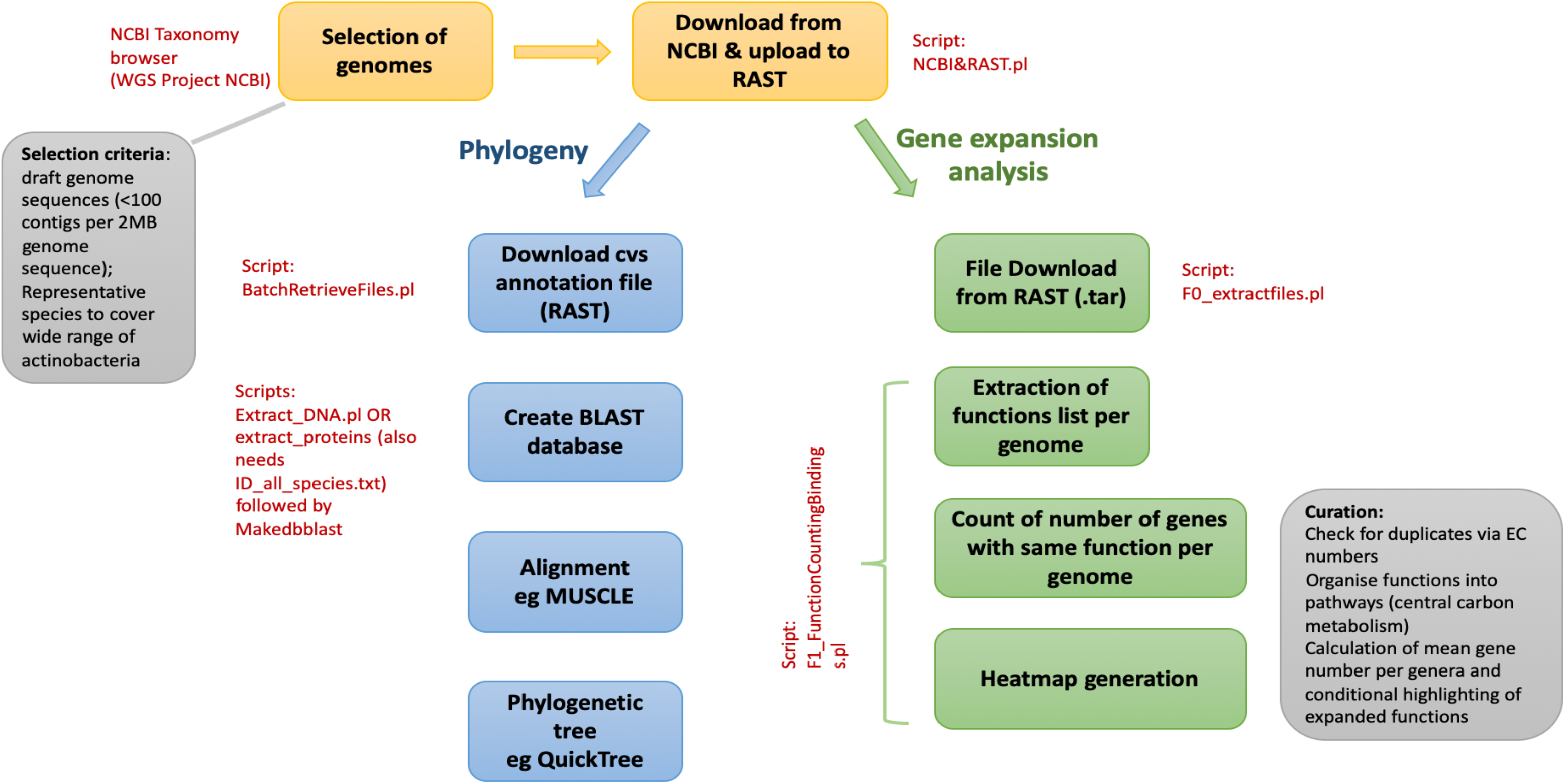
Schematic workflow for the creation of the ActDES database. Genomes were selected from NCBI Taxonomy browser and uploaded for annotation to RAST[38]. The annotated genomes were the processed for two different analyses. Firstly, the functional roles were downloaded and for each functional role the numbers of occurrences per genome were counted in order to obtain an expansion table (**Supp. Table S2**) by comparing the mean of each genus to the overall mean of all genera. Secondly, the genomes were used to extract all nucleotide and protein sequences in FASTA format which could then be queried by sequence using BLAST [39]. The hits were aligned in MUSCLE [16] and after refinement the alignment was used to construct phylogenetic trees in Quicktree [17].

The database was generated via the NCBI taxonomy browser (https://www.ncbi.nlm.nih.gov/Taxonomy/Browser/www.tax.cgi) to identify Actinobacterial genome sequences. The quality of the genome sequences was filtered by the number of contigs (<100 contigs per 2Mb of genome sequence) and the genomes were downloaded from the NCBI WGS repository (https://www.ncbi.nlm.nih.gov/Traces/wgs/). These genomes were then dereplicated to ensure that the database comprised a wide taxonomic range of the phylum, resulting in 612 species from 80 genera within 13 suborders of the Actinobacteria (**Supp. Table S1**).

Each of these 612 genomes was reannotated using RAST. Default settings were used to ensure equivalence of annotation across the database and the annotation files of each genome were downloaded (Data File: cvs files). These annotation files were subsequently used to extract all protein and nucleotide sequences into two files. Each of these files was subsequently converted into BLAST databases (a protein database and a nucleotide database https://doi.org/10.6084/m9.figshare.12167724) to facilitate phylogenetic analysis. Sequences of interest can be aligned using MUSCLE [16] and phylogenetic trees constructed using a range of tree construction software such as QuickTree [17], IQ tree [18] or MrBayes [19]. Subsequent trees may be visualized in software such as FigTree v1.4.2 (http://tree.bio.ed.ac.uk/software/figtree/).

The RAST annotation files were also used to extract the functional roles of each CDS per genome and the level of gene expansion was assessed for each genome by counting the number of genes per species per functional category (gene function annotation). The dataset was then curated manually for central carbon metabolism and amino acid biosynthesis pathways to create the gene expansion table (**Supp. Table S2**) with the organisms grouped according to their taxonomic position. The quality of the data was checked at each step for duplicates and inconsistencies and was curated manually to exclude faulty entries.

As the NCBI taxonomy browser database is overrepresented in *Streptomyces* genomes, this is also reflected in the ActDES database (288 *Streptomyces* genomes from a total of 612 genomes). However this was addressed in the expansion table (**Supp. Table S2**) by calculating the mean occurrence of each functional category within each genus and then calculating an overall mean for the phylum to compensate. The mean occurrence of each functional category per genus plus the standard deviation was also calculated and this was used to analyse the occurrence of each functional gene category per species within **Supp. Table S2**. A gene function annotation with a gene copy number value above the mean plus the standard deviation for each genus, indicated that there had been a gene expansion event in that species and this was noted. The gene expansion table (**Supp. Table S2**) enables researchers to identify groups of genes of interest for subsequent phylogenetic and evolutionary analysis, which can be performed with confidence due to the highly curated nature of the data included in the database.

## Results

The gene expansion table (**Supp. Table S2**) lists 612 species of 80 genera within the Actinobacteria with data that provides an extensive analysis at the phylum level, which is the starting point for detailed phylogenomic studies. Gene expansions were identified in separate datasets at the genus and species levels, along with details of the numbers of genes in each functional category per species and the average numbers of genes in each functional category per genus expanded within the genomes. These data can be used subsequently in phylogenomic analyses to identify targets for metabolic engineering and gene function studies. Identification of expanded gene families may also facilitate the recognition of novel natural product biosynthetic gene clusters, for which gene expansion events of primary metabolic genes have been classified to be associated within BGCs as biosynthetic enzymes or through provision of additional copies of antibiotic targets that may subsequently function as resistance mechanisms [6, 11, 20–24].

This database has found utility for studying primary metabolic gene expansions in *Streptomyces*. It enabled a detailed *in silico* analysis of the duplication event leading to the two pyruvate kinases in the genus of *Streptomyces* subsequently enabling the functional characterisation of the two isoenzymes to reveal how they contribute to metabolic robustness [15]. ActDES may also be useful for investigating the distribution of primary metabolic genes across the phylum to link phenotype to genotype and phylogenetic position. An initial RpoB phylogeny has been constructed previously using this database [15] which provided a robust universal phylogeny for comparison of individual protein trees[25].

To demonstrate the utility of ActDES, the glucose permease/glucokinase system of the Actinobacteria was investigated. The role of nutrient-sensing in regulation of antibiotic biosynthesis is well known [26] with the enzyme glucokinase (Glk) playing a central role in carbon-catabolite repression in *Streptomyces* [27]. In most bacteria, CCR is mediated by the phosphoenolpyruvate-dependent phosphotransferase system (PTS), yet in *Streptomyces*, glucose uptake is mediated by the Major-facilitator superfamily (MFS) transporter, glucose permease (GlcP), and there is evidence for direct interaction between Glk and GlcP which may mediate CCR [28]. Understanding the nature and distribution of these enzymes will play a key role in developing industrial fermentations with glucose as major carbon source. Investigating the distribution of the glucose permease/glucokinase system across the phylum shows that GlcP and Glk have been the subject of gene expansion events in some members of the Streptomycetales, most notably the *Streptomyces*, with a patchy distribution of the Glk/GlcP system across the remainder of the phylum (**Table S2; Genus tab**). However, where the Glk/GlcP system is found, the number of expansion events observed is greater for Glk than for GlcP (**Fig. 2 A & B**). The phylogenetic trees (**Fig. 2A & 2B**) clearly show two clades for Glk and GlcP within the Streptomycetales (Interactive trees are available via Microreact [29]: Glk https://microreact.org/project/w_KDfn1xA/90e6759e and GlcP https://microreact.org/project/VBUdiQ5_k/0fc4622b). However, these clades differ in the number of sequences, with the Glk clades being equal in number, suggesting that a duplication event has occurred within the Streptomycetales (**Fig. 2A**). The is consistent throughout the order, with the patterns largely the same as that observed for *S. coelicolor*. This species has two ROK-family ATP-dependent glucokinases, SCO2126 (*glkA*) and SCO6260, that share around 50% amino acid sequence identity and each is found in one of the distinct clades (permease-associated kinases and orphan kinases; **Fig. 2A**). Whilst SCO2126 is a GlcP-associated kinase, the gene encoding SCO6260 is located in an operon including genes encoding a putative carbohydrate ABC-transporter system, which has been reported previously [30]. SCO6260 appears to be the only glucokinase in the database that is associated with an ABC-transporter. This may suggest that expansion of the Glk gene family in Streptomycetales might have occurred to extend the number of CCR-mediating kinases in the genome, adding increased regulatory complexity to carbohydrate metabolism in this group of organisms that use CCR as a major regulator of specialised metabolism.

**Fig. 2.**
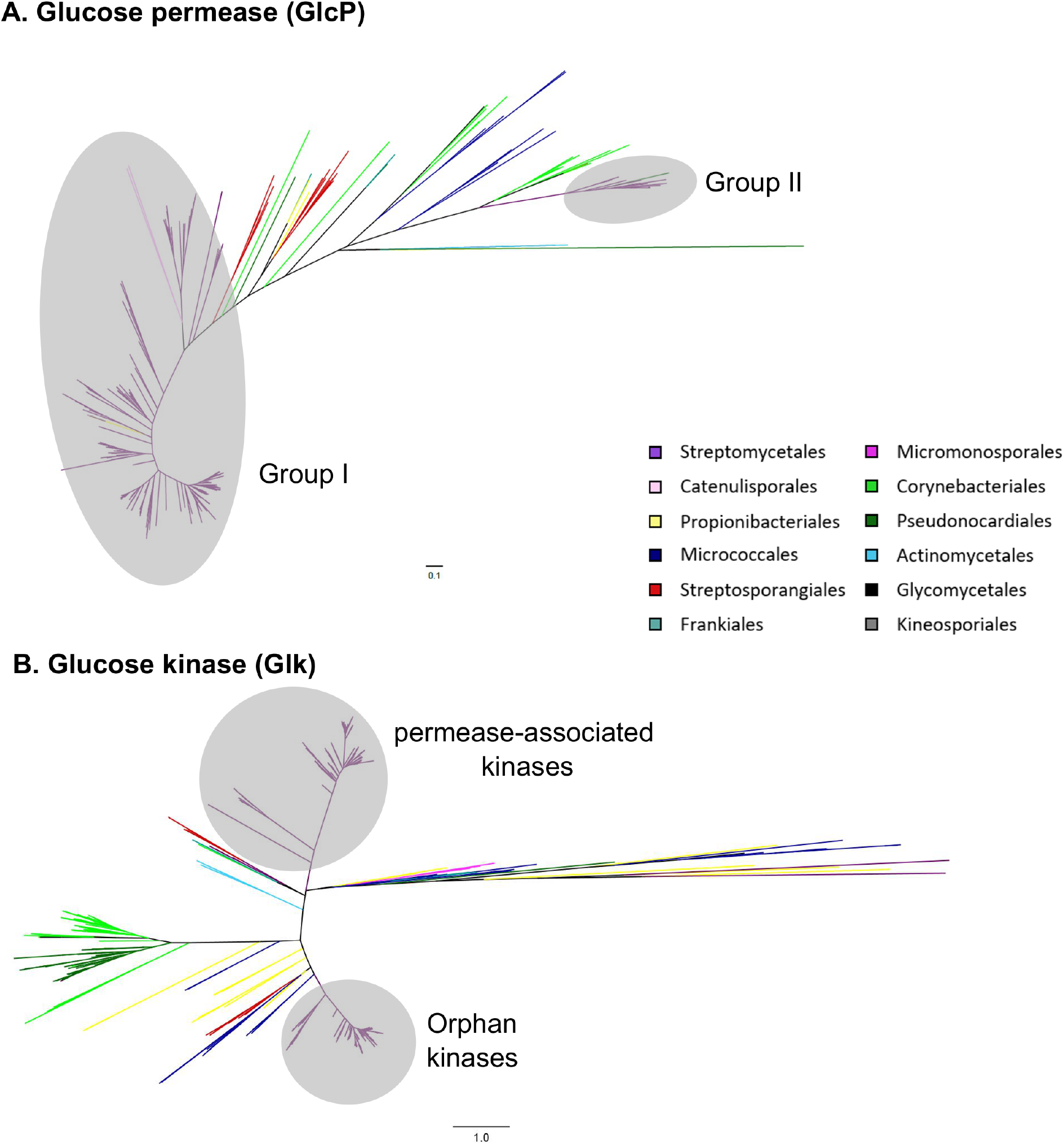
**A)** Actinobacterial-wide phylogenetic tree of glucose permeases (GlcP) **B)** Actinobacterial-wide phylogenetic tree of Glucokinases (Glk). Trees are colour-coded according to the NCBI Taxonomy browser (https://www.ncbi.nlm.nih.gov/Taxonomy/Browser/www.tax.cgi). Interactive trees are also available via Microreact [29]: GlcP https://microreact.org/project/VBUdiQ5_k/0fc4622b) and Glk https://microreact.org/project/w_KDfn1xA/90e6759e.

The two clades for GlcP within the Streptomycetales differ in size suggesting either gene duplication followed by gene loss, or an expansion through horizontal gene transfer (HGT) has occurred. A detailed examination of these clades by species (**Table S2; Species Tab**) shows the presence of both scenarios. There are duplicated enzymes located within the same clade (as observed in *S. coelicolor;* Group I) or additional copies of the permease which are located in a phylogenetically distinct clade, which remarkably consists entirely of sequences from the genus *Streptomyces* (Group II; **Fig. 2B**). This suggests that they may have been acquired via HGT. The expansive nature of the duplicated Glk enzymes compared to GlcP may be due to the role played in CCR by the GlkA enzymes [27] and the different transcriptional activities under glycolytic and gluconeogenic conditions [31], yet quite how these different Glk enzymes interact with the permease(s) under various conditions requires further experimental investigation to understand their exact physiological role, and how this may be translated in to industrial strain improvement processes.

## Discussion

Large scale WGS and phylogenomic analysis is increasingly used for identifying targets for genome and metabolic engineering, studies of metabolic capabilities, pathogen phylogenomics and evolutionary studies. These studies are often complicated by the large number of sequences in the databases, database redundancy and the poor quality of some genome sequence data. The development of the high-quality, curated ActDES database, reported here, enables phylum-wide taxonomic representation of the Actinobacteria coupled with quality-filtered genome data and equivalent annotation for each CDS.

The intended primary use of ActDES will be in the study of primary metabolism, but it is not limited. It can also inform the development and evolution of metabolism in strains that produce bioactive metabolites, given the high representation of genera renowned for their ability to produce natural products such as *Streptomyces* and *Micromonospora*. Due to a greater understanding of BGC evolution and genome organisation in Actinobacteria it is becoming increasingly clear that genes whose functions are in primary metabolism may actually contribute directly to the biosynthesis of specialised metabolites and, hence, the identification of duplicates may indicate the presence of cryptic BGCs [6, 11] or, when associated with precursor biosynthetic genes, provide the raw material for the enzymes across multiple BGCs [32–34].

ActDES may also find utility in evolutionary studies of expanded gene families across the Actinobacterial phylum that contribute to virulence such as the *mce* locus which is known to facilitate host survival in Mycobacteria[35], but also facilitates xenobiotic substrate uptake in *Rhodococcus* [36] and enables root colonization and survival in *Streptomyces* [37]. With phylum-wide taxonomic representation of established Actinobacterial animal and plant pathogens, the scope for evolutionary studies using these data is enormous.

## Usaqe Notes

The CVS files of each genome contains the RAST annotation details in addition to the DNA and protein sequences for each annotated CDS (https://doi.org/10.6084/m9.figshare.12167880). The Genome list contains the RAST ID (which is equivalent to the name of the .cvs file) along with the NCBI ID (sequence ID; **Table S1**) plus the species name which are included in the dataset. Further details of annotating batches of genomes in RAST can be found here https://github.com/nselem/myrast.

The primary metabolism expansion tables (**Supp. Table S2**) are organised by metabolic pathway along the top row with the Enzyme Commission (EC) number and functional annotation, with the first column being the taxonomic assignment. The genus table shows the average number of genes of the annotated function. Highlighted cells reflect gene expansion events, i.e. those genes that are present in higher number than the overall mean across the database plus the standard deviation.

It is suggested that the gene expansion table (**Supp. Table S2**) is searched in the first instance (either by species or genus of interest or by a specific enzymatic function). This can be carried out by a simple text search. This will then allow the identification of a query sequence from a species or gene of interest (either nucleotide or amino acid sequence) which can then be searched against the curated BLAST database allowing a detailed phylogenetic analysis of a gene/protein of interest by using standard alignment and tree building software tools. These data can also be used in detailed evolutionary analysis of selection, mutation rates etc.

## Author Contributions

Conceptualization: JKS, NS-M, PAH, FB-G

Data curation: JKS & NS-M

Formal analysis: JKS, N.S-M, PC-M, & ASB

Funding acquisition: PAH, FB-G

Methodology: JKS, NS-M, PC-M, FB-G & PAH

Project administration: PAH & FB-G

Supervision: PAH, ISH & F.B-G

Writing – original draft: JKS, ASB & PAH

Writing – review and editing: JKS, N.S-M, PC-M, ASB, ISH, FB-G & PAH

## Acknowledgements

We thank the Scottish Universities Life Science Alliance (SULSA) for BioSkape PhD funding to JKS, and a Mac Robertson Travelling Scholarship awarded to JKS to visit the laboratory of FB-G., an Industrial Biotechnology Innovation Centre (IBioIC) and GlaxoSmithKline funded PhD studentship to ASB, and a NERC (grant NE/M001415/1), BBSRC (grants BB/N023544/1 and BB/T001038/1) and BBSRC/NPRONET (grant NPRONET POC045) to PAH. Work in the laboratory of FB-G. was funded by CONACyT, Mexico Metabolic Robustness in Streptomyces, and Langebio institutional funds to support PC-M. as a postdoctoral fellow and Royal Society Newton Advanced Fellowship (NAF\R2\18063).

## Ethical statement

No ethical approval was required.

